# Arterio-Venous Remodeling in the Zebrafish Trunk is Controlled by Genetic Programming and Flow-Mediated Fine-Tuning

**DOI:** 10.1101/403550

**Authors:** Ilse Geudens, Baptiste Coxam, Silvanus Alt, Véronique Gebala, Anne-Clémence Vion, Andre Rosa, Holger Gerhardt

## Abstract

How developing vascular networks acquire the right balance of arteries, veins and lymphatics to efficiently supply and drain tissues is poorly understood [1, 2]. In zebrafish embryos, the robust and regular 50:50 global balance of intersegmental veins and arteries that form along the trunk [3], prompts the intriguing question how the organism keeps “count”. Previous studies suggest that the ultimate fate of an intersegmental vessel (ISV) is determined by the identity of the approaching secondary sprout emerging from the posterior cardinal vein (PCV) [1, 4-7]. Here, using high time-resolution imaging, advanced cell tracking and computational analysis, we show that the formation of a balanced trunk vasculature involves an early heterogeneity in endothelial cell (EC) behavior in the seemingly identical primary ISVs and an adaptive flow-mediated mechanism that fine-tunes the balance of arteries and veins along the trunk. Detailed examination of the trunk vasculature dynamics throughout development reveals the frequent formation of three-way vascular connections between primary ISVs, the dorsal aorta (DA) and the PCV. Differential resolution of these connections into arteries or veins is mediated by polarized cell movement of the ECs within the ISV. Quantitative analysis of the cellular organization, polarity and directional movement of ECs in primary ISVs identifies an early differential behavior between future arteries and veins that is largely specified in the ECs of the individual ISVs, is dependent on Dll4/Notch, and occurs even in the absence of secondary sprouting. Notch signaling is involved in a local patterning mechanism normally favoring the formation of alternating arteries and veins. The global artery-vein balance is however maintained through a flow-dependent mechanism that can overwrite the local patterning. We propose that this dual mechanism driving arterio-venous identity during developmental angiogenesis in the zebrafish trunk provides the adaptability required to establish a balanced network of arteries, veins and lymphatic vessels.

---

The zebrafish trunk vasculature first arises as an all-arterial network and is subsequently remodeled into an efficiently perfused network of arteries and veins. Although the order of arteries and veins along the trunk is not fixed, every embryo forms a balanced number of arteries and veins (Figure 1A) [3]. How exactly this remodeling is organized to result in this balance is unknown. Current concepts favor the idea that secondary sprouts are fate-restricted by their level of expression of the lymphatic determinant Prox1; cells expressing low levels of Prox1 will connect to the primary ISV and form a vein, whereas sprouts expressing high levels continue to the level of the horizontal myoseptum and give rise to the lymphatic precursor structures [5, 6].

**Figure 1.**
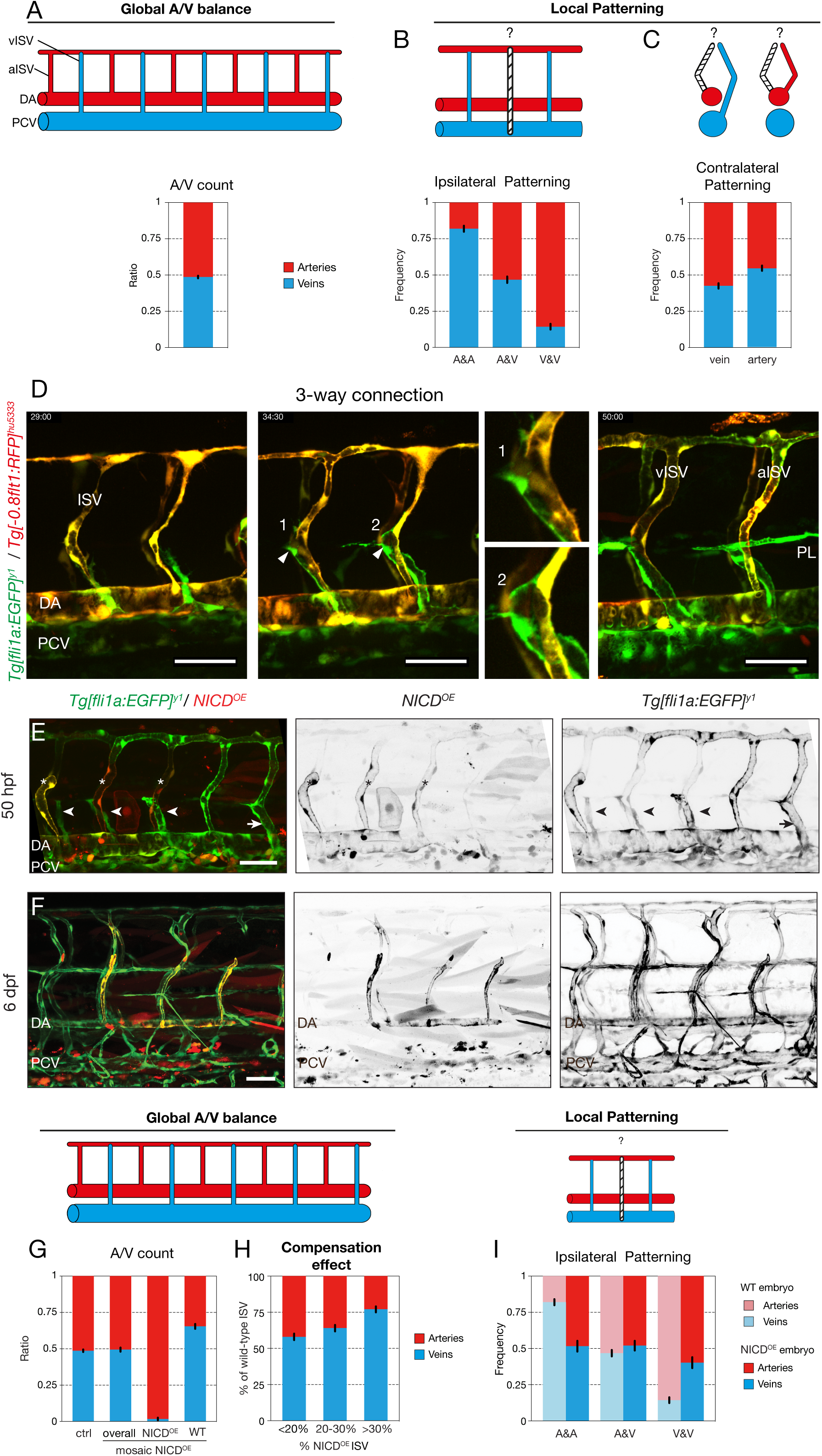
Notch mediates the local patterning of the trunk vasculature. A) Quantification of the ratio of arterial and venous ISVs in a 10-somites region of the trunk of 6 dpf WT embryos (n=3 experiments, 74 embryos). B) Ipsilateral neighborhood analysis of vessel identity with 2 neighbors in 6 dpf WT embryos (n=3 experiments, 74 embryos, 1184 ISVs). C) Contralateral neighborhood analysis of vessel identity in 6 dpf WT embryos (n=3 experiments, 74 embryos, 1480 ISVs). D) Stills from time-lapse movie (Supplementary Movie S1) of a *Tg[fli1a:EGFP]*^*y1*^ */ Tg[-0.8flt1:RFP]*^*hu5333*^ embryo showing ISV remodeling into an aISV and a vISV. In both cases, a lumenized connection is formed between the secondary sprout and the primary ISV (arrowhead). In the case of the formation of an aISV, the connection is lost again and the secondary sprouts forms lymphatic precursors at the horizontal myoseptum (parachordal lymphangioblasts, PL). In case of vISV remodeling, the secondary sprout connection is stabilized and the connection between primary ISV and DA regresses. E) *Tg[fli1a:EGFP]*^*y1*^ embryos mosaically expressing a pT2Fli1ep-zN1aICD-basfli-mCherry construct at 50 hpf. Lymphangiogenic sprouts i.e. sprouts delivering lymphatic precursors at the horizontal myoseptum (arrowheads), can be observed at the position of NICD overexpressing (NICD^OE^) ISVs (asterisks) F) *Tg[fli1a:EGFP]*^*y1*^ embryos mosaically expressing a pT2Fli1ep-zN1aICD-basfli-mCherry construct at 6dpf. NICD^OE^ mCherry-positive cells were found almost exclusively in the arterial compartment of the vasculature. G) Quantification of the ratio of arterial and venous ISVs in a 10-somites trunk region of 6dpf control embryos (n=3 experiments, 74 embryos) and mosaic NICD^OE^ embryos (n=3 experiments, 51 embryos). In mosaic embryos the arterio-venous distribution was quantified overall and separately in NICD^OE^ and wild-type ISVs. Wild-type ISVs compensate for the forced arterialization of NICD^OE^ ISVs by increasingly being specified into vISVs. H) Quantification of the percentile presence of arterial and venous ISVs in a 10-somites region of the trunk of 6dpf NICD^OE^ embryos. Mosaic NICD^OE^ embryos represented in panel G were grouped based on their relative number of NICD^OE^ ISVs (<20, 20-30 or >30%; n=13, 26 and 12 embryos, respectively). I) Ipsilateral neighborhood analysis of vessel identity with 2 neighbors in 6dpf NICD^OE^ embryos (n=3 experiments, 40 embryos, 592 ISVs) compared to WT embryos (n=3 experiments, 74 embryos, 1184 ISVs). DA, dorsal aorta; PCV, posterior cardinal vein; ISV, intersegmental vessel; PL, parachordal lymphangioblasts; WT, wild-type; NICD^OE^, NICD-overexpressing. Scale bars, 50μm

To explore the nature of the global artery-vein balance, we analyzed the sequence of arteries and veins on both sides of a 10-somite segment in 6 days post fertilization (dpf) wild-type embryos, in which arteries and veins are already functionally specified, and ran a neighborhood analysis to determine the conditional probabilities of forming an artery or a vein given the status of the neighboring vessels. This showed a strong, albeit imperfect ipsilateral patterning of alternating vessel fates (Figure 1B, Figure S1A-B). A similar analysis revealed a weak contra-lateral patterning between opposing vessels in the zebrafish trunk, although it is only a weak predictor of vessel identity (Figure 1C). Consequently, an ISV surrounded by arterial ISVs (aISV) has a strong probability of being a venous ISV (vISV).

Previous studies proposed that the initial event driving patterning occurs in the PCV, establishing either venous or lymphatic cell fate in cells forming the secondary sprouts, thereby dictating whether they connect to the primary ISV to form a vein, or not, thus forming lymphatics. Surprisingly, we instead find regular interactions between the primary ISV and the secondary sprout irrespective of the final outcome of the patterning event (Figure 1D; Supplementary Movie S1). In most segments, secondary sprouts fuse with the primary ISV, forming three-way connections between the DA, ISV and PCV. Ultimately however, the connection to the DA either regresses, turning the ISV into a vein, or remains stable, thus preserving arterial identity while the secondary sprout disconnects from the ISV and contributes to lymphatic formation (Figure 1D). Occasional occurrence of these three-way connections has previously been reported, albeit in only 3% of the ISVs [2]. Interestingly, our quantification reveals that at least 77.5% of future aISVs (N = 40) are transiently connected to secondary sprouts, forming a lumenized and perfused shunt (Supplementary Figure S1C, Supplementary Movie S2).

Because of the known role of Notch in regulating arterio-venous specification [8-11], and the increased formation of vISVs upon Notch inhibition [1, 12], we asked whether Notch activity cell-autonomously influences ISV specification. We used Tol2 transgenesis to mosaically overexpress the intracellular domain of Notch (NICD) together with a mCherry reporter in single ECs of *Tg[fli1a:EGFP]*^*y1*^ embryos. mCherry-positive NICD-overexpressing cells were found almost exclusively in the arterial compartment of the trunk vasculature (DA and aISVs) (Figure 1E-G), indicating that Notch signaling might play a cell-autonomous role in ISV patterning. Live imaging revealed that the ISV containing NICD-overexpressing cells remained arterial, while still forming transient lumenized three-way connections (Supplementary Figure S1D, Supplementary movie S3). Interestingly, when studying all vessels in these embryos, we observed a strong bias towards the formation of vISVs in ISVs containing only WT cells (Figure 1G). Moreover, when analyzing individual embryos, we observed that the more ISVs contained NICD-overexpressing cells, the more wild-type ISVs turned into veins (Figure 1H). As a result, the global artery to vein balance was maintained in embryos with mosaic overexpression of NICD (Figure 1G). Neighborhood analysis in these embryos showed a complete disruption of ipsilateral patterning, rendering all local patterns equally probable (Figure 1I). Taken together, these results suggest the existence of a compensation mechanism that maintains the global balance between arteries and veins, and which operates in addition to or independent of the local patterning.

Since a balanced artery-vein network is required for optimal blood flow distribution in the fish trunk, we speculated that flow and flow sensing play an important role in the patterning and/or compensation of vessel specification. To test this hypothesis, we slowed down the heart rate by treating embryos with tricaine (tricaine mesylate, MS-222), a muscle relaxant commonly used as an anesthetic in fish. Treatment of the embryos with twice the dose normally used for anesthesia, after the onset of secondary sprouting, from 31 to 52 hpf, significantly reduced blood flow (heart rate 80.9% ± 10.8 of WT). Flow reduction in wild-type embryos disrupted the global balance of arteries and veins at 6dpf (61.2% ± 0.06 aISV) (Figure 2A) whilst ipsilateral patterning was widely retained (Figure 2B), indicating that blood flow could be a critical determinant in the compensation mechanism that establishes the overall balance in the number of arteries and veins along the trunk. Indeed, treating embryos harboring mosaic overexpression of NICD in the vasculature with a similar dose of tricaine resulted in a reduction in the number of WT vessels becoming vISVs, thus abolishing the compensation effect (Figure 2C).

**Figure 2.**
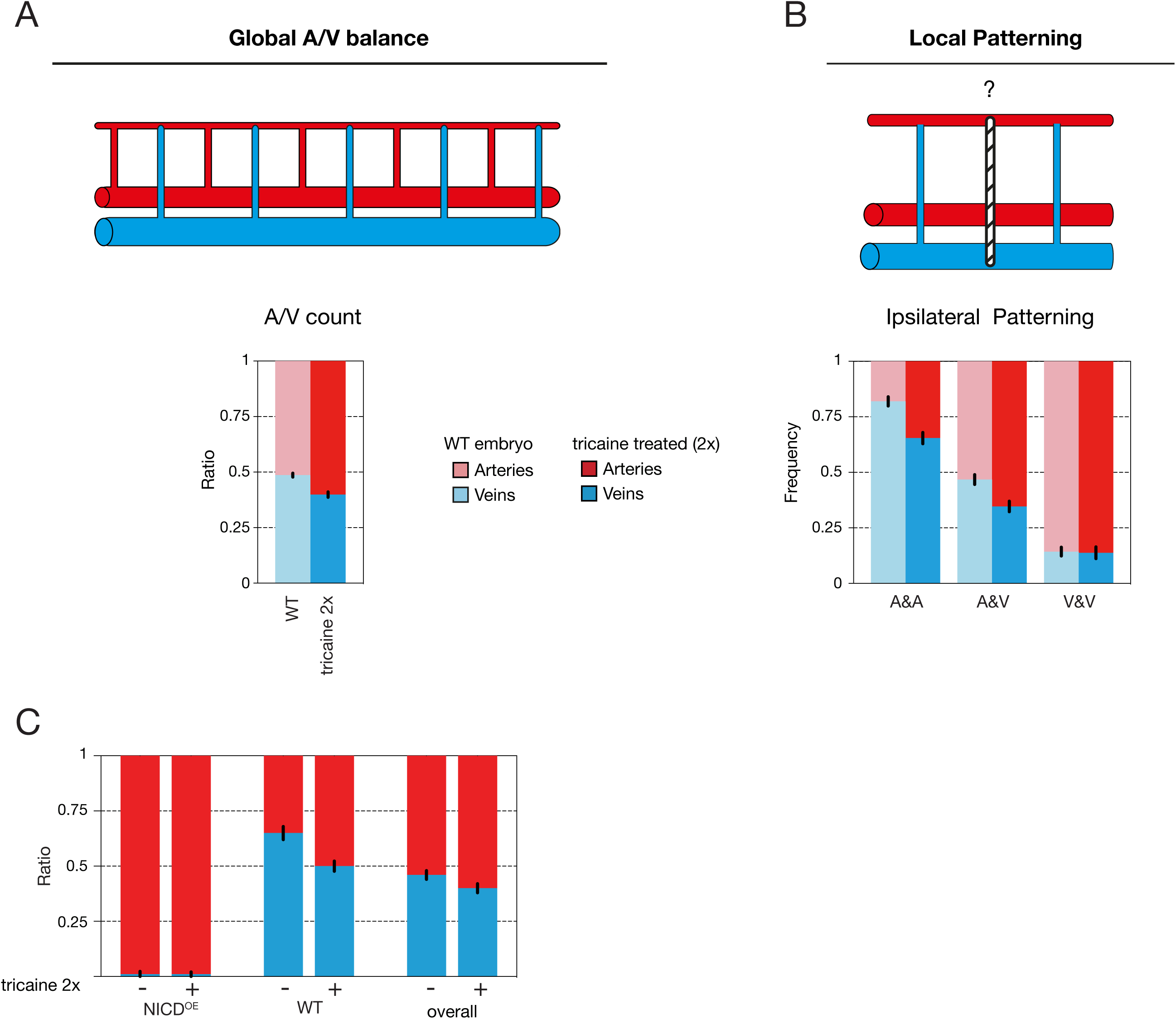
Flow mediates the global arterio-venous balance of the trunk vasculature. A) Quantification of the ratio of arterial and venous ISVs in a 10-somite region in the trunk of 6 dpf wild-type embryos either untreated (control; n=3 experiments, 74 embryos) or treated with 2x tricaine to slow down the heart rate (n=3 experiments, 65 embryos). B) Ipsilateral neighborhood analysis of vessel identity with 2 neighbors in 6 dpf wild-type embryos either untreated (control; n=3 experiments, 74 embryos, 1184 ISVs) or treated with tricaine to slow down the heart rate (n=3 experiments, 65 embryos, 950 ISVs). C) Quantification of the ratio of arterial and venous ISVs in a 10-somite trunk region of 6 dpf mosaic NICD^OE^ embryos either untreated (-) or treated (+) with 2x tricaine to slow down the heart rate (respectively: n=3 experiments, 51 embryos; n=2 experiments, 20 embryos). In mosaic embryos the arterio-venous distribution was quantified overall and separately in mosaic NICD^OE^ and wild-type ISVs (WT). Blood flow reduction eliminates the compensating venous remodeling in wild-type (WT) ISVs in mosaic NICD^OE^ embryos, resulting in and overall shift towards more arterial ISVs.

The question of which of the branches of the three-way connection is stabilized or lost is reminiscent of the pruning process in capillary networks during flow-dependent remodeling in mouse retina and zebrafish brain [13, 14]. Previous studies of developmental vessel pruning established that ECs exposed to physiologically high levels of blood flow, above a certain threshold, align in the direction of the flow and polarize against it [14, 15]. As a consequence, adjacent vessel segments (branches) that experience different levels of flow show opposite endothelial polarization and movement. ECs polarize and migrate away from the branch experiencing low or sub-threshold flow, into the branch under high flow. The divergent polarity of cells in the low-flow branch causes cells to disconnect. Therefore, we analyzed the polarity of ECs during the remodeling process in *Tg[fli1a:GFP]*^*y1*^*;Tg[fli1a:B4GalT-mCherry]*^*bns9*^ embryos during three different phases: I) before secondary sprout connection, II) when the three-way connection is present and III) after resolution (Figure 3A). As expected, we found that most ECs polarize against the flow, leading to ventral polarity in aISVs and dorsal polarity in vISVs (Figure 3B-E, Supplementary movies S4-S5). Surprisingly, ECs in future aISVs and vISVs already showed differential polarization patterns before secondary sprout connection (Figure 3D,E - phase I). Moreover, when tracking the nucleus of ECs in *Tg[- 0.8flt1:RFP]*^*hu5333*^;*Tg[fli1a:eGFP]*^*y1*^ embryos in all three phases, we observed that cells constituting the primary ISV exhibited dorsal-oriented movement in future vISVs, whereas cells in future aISVs remained largely at the same position or moved slightly ventrally (Figure 3F). Again, this difference was already present in phase I, suggesting that specification of arteries and veins occurs within the primary ISVs prior to interactions with the secondary sprouts from the PCV.

**Figure 3.**
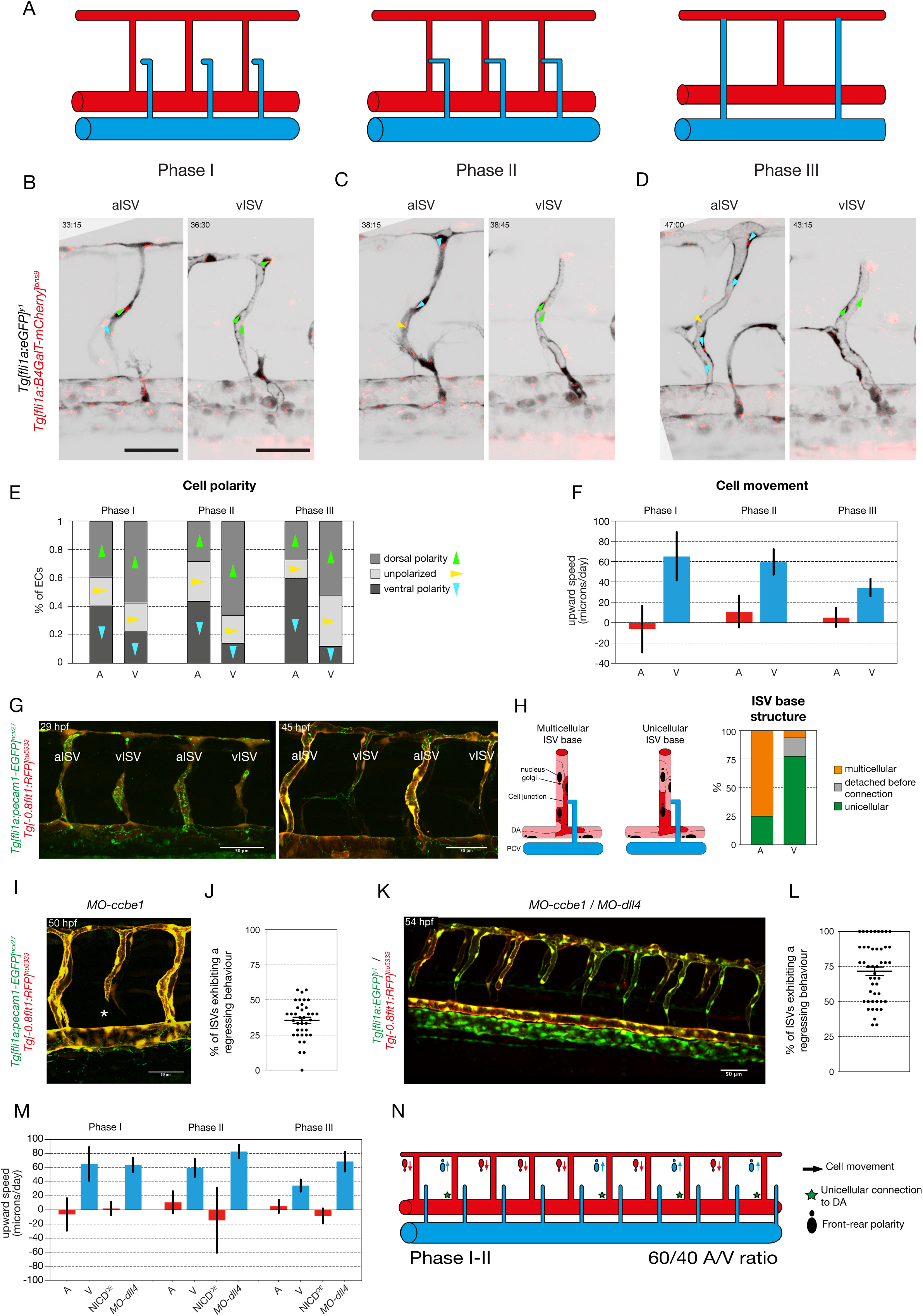
primary ISVs are specified into aISVs and vISVs prior to connection to secondary sprouts originating from the PCV. A) Schematic representation of the three phases of primary ISV remodeling; phase I: before secondary sprout connection to the primary ISV, phase II: When a lumenized connection is formed between the secondary sprout and the primary ISV, phase III: when the three-way connection resolves into aISV or vISV. B-D) Stills from time-lapse movies (see Supplementary Movie S4-S5) of ECs polarity in aISVs and vISVs of Tg*[fli1a:GFP]*^*y1*^*;Tg[fli1a:B4GalT-mCherry]*^*bns9*^ embryos during the three different phases: I) before secondary sprout connection, II) during three-way connection and III) after resolution. Arrows point from the center of the nucleus to the center of the golgi complex: green arrows indicate dorsal polarity, blue arrows indicate ventral polarity, yellow arrows indicate unpolarized ECs. E) Quantification of EC polarity in aISVs (n=7 aISVs, 16 cells) and vISVs (n=8 vISVs, 17 cells) of *Tg[fli1a:GFP]*^*y1*^*;Tg[fli1a:B4GalT-mCherry]*^*bns9*^ embryos during the three different phases: I) 2.5 hours before secondary sprout connection, II) during three-way connection and III) 2.5 hours after resolution. F) Quantification of ECs upward speed (in microns/day) in aISVs (n= 12 aISVs, 67 cells) and vISVs (n=13 vISVs, 103 cells) during the three different phases: I) 1 hour before secondary sprout connection, II) during three-way connection and III) 1 hour after resolution. G) Stills from time-lapse movie (Supplementary Movie S6) of a *Tg[fli1a:pecam1-EGFP]*^*ncv27*^*; Tg[-0.8flt1:RFP]*^*hu5333*^ embryo showing ISV remodeling into aISVs and vISVs at 29 and 45 hpf. H) Quantification of the cellular structure at the base of the primary ISV at the inception of Phase II. Detection or absence of Pecam1-eGFP junctional expression is used to characterize the nature of the connection (unicellular or multicellular) (n=5 experiments, 34 embryos, 12 aISVs, 40 vISVs). I) Still from time lapse movie (see supplementary movie S7) of a T*g[fli1a:pecam1-EGFP]*^*ncv27*^ *; Tg[-0.8flt1:RFP]*^*hu5333*^ 5ng *MO-ccbe1* embryo showing ISV regression in the absence of secondary sprouting. J) Quantification of percentage of primary ISV exhibiting a regression behavior (full disconnection from the DA, thin membrane connection to the DA, lumen collapse and reconnection, and cell death at the base of the primary ISV – see Supplementary Figure 2B) (n=4 experiments, 37 morphants, 29 WT controls, 241 morphant vessels). K) Stills from time-lapse movie (Supplementary Movie S8) of a *Tg[fli1a:EGFP]*^*y1*^ */ Tg[- 0.8flt1:RFP]*^*hu5333*^ *5ng MO-ccbe1 / 10ng MO-dll4* embryo showing ISV regression in the absence of secondary sprouting. L) Quantification of percentage of primary ISV exhibiting a regression behavior (full disconnection from the DA, thin membrane connection to the DA, lumen collapse and reconnection, and cell death at the base of the primary ISV – see Supplementary Figure 2C) (n=4 experiments, 47 morphants, 17 WT, 402 morphant vessels). M) Quantification of ECs upward speed (in microns/day) in ISVs of WT (n=12 aISV (67 cells), 13 vISV (103 cells)), NICD^OE^ (n=30 aISV, 29 NICD^OE^ cells) and *MO-dll4* (n=9 vISV, 85 cells) embryos (32 to 54 hpf) at 3 different time points: I) before secondary sprout connection, II) during three-way connection and III) after resolution. N) Schematic representation of ISV specification prior to and at the inception of the three-way connection, quantifiable through EC polarity, upward movement speed and cellular structure at the connection to the DA. Scale bars, 50μm

Since vessels regression events are characterized by a progressive conversion from multicellular to unicellular arrangements [16], we analyzed the remodeling process at the cellular level by imaging cellular junctions at the base of the primary ISV in *Tg[fli1a:pecam1-EGFP]*^*ncv27*^*;Tg[-0.8flt1:RFP]*^*hu5333*^ embryos during remodeling. For most future vISVs, at the moment of connection to the secondary sprout, the base of the ISV was made of a single EC connecting the vessel to the DA, whereas the majority of future aISVs had a multicellular arrangement at the ISV base (Figure 3G-H, Supplementary movie S6). This suggests that the observed cellular behavior within the primary ISV, which drives stabilization or regression of the DA connection, is already structurally specified before the time of connection to the secondary sprout. Combined, these findings uncover a heterogeneity in primary ISVs, showing differential behavior of ECs that is predictive of their specification, prior to the connection to venous derived secondary sprouts (Figure 3N). These findings also suggest the possibility that the process of disconnecting from the DA to form a vISV may be initiated independently of the approaching secondary sprout.

To test this hypothesis, we inhibited the formation of secondary sprouts by inactivating *ccbe1*, a critical mediator of Vegfc processing [17]. Despite the absence of secondary sprouts in *ccbe1* morpholino treated embryos, a subset of ISVs (on average 38.2% ± 4.8) showed a dynamic behavior consistent with regression of the DA connection, indicating that vein-specific behavior of ISV ECs proceeds even in the absence of secondary sprouts (Figure 3I-J, Supplementary movie S7). Regressing behavior was evident by the presence of only a thin membrane connection, lumen collapse and reconnection, and even full detachment of the ISV from the DA in 36.6% (± 11.32) of regressing ISVs (Supplementary Figure S2A-B). Interestingly, co-injection with a *dll4* morpholino, which leads to the formation of an increased number of veins in the trunk [18], resulted in a dramatic increase of ISV regression (*MO-dll4* + *MO-ccbe1*: 71.6%; ± 21.1) (Figure 3K-L, supplementary Figure S2C, Supplementary movie S8), strongly suggesting that the autonomous regression behavior observed in the absence of secondary sprouts is associated with a venous specification that is established early on in primary ISVs.

Given the strong influence of Notch activity on primary ISV specification, we analyzed the effect of manipulating Notch signaling on EC behavior within the ISV. Tracking of cell movement in *dll4* morphants showed that the primary ISV cells migrated dorsally like wild-type venous cells, while in ISV overexpressing NICD, ECs instead remained at the same position or migrated ventrally like arterial cells. Both behaviors are visible in all phases of remodeling (Figure 3M). This indicates that Notch has an effect on vessel specification prior to the connection with a secondary sprout. Analysis of cell polarity during remodeling in *dll4* morphant embryos showed that the majority of cells are polarized dorsally, as in wild-type vISVs, but that an increased number of cells appear unpolarized (polarized orthogonally to the ISV) (Supplementary Figure S2D). Similarly, in flow chamber experiments under shear stress conditions mimicking physiological flow, we found that chemical inhibition of Notch signaling using the gamma-secretase inhibitor DAPT prevented HUVECs from efficiently orienting parallel to the flow and from polarizing against the flow (Supplementary Figure S2E).

Thus, our combined *in vivo* and *in vitro* results indicate that Notch signaling influences EC polarity and movement. Interestingly, the polarization of ECs in primary ISVs in phase I is similar to that later observed in functionally specified ECs polarizing against flow [14, 15, 19], suggesting that Notch- and flow-mediated polarization could be established through similar cellular mechanisms.

Similar to vascular regression processes in the zebrafish brain and eye vasculature, our work indicates that the pruning of transient arterio-venous three-way connections that resolve into aISV or vISV is dependent on directional movement of the EC [13, 20]. When looking at the direction of EC movement we can distinguish between ISVs with dorsal and ventral movement. In the first case, divergent polarity between the ECs in the ISV and in the secondary sprout connection will favor disconnection of the PCV branch. In the second case, dorsal polarization of ECs that connect the ISV to the DA will favor their disconnection from the DA, whilst converging with the polarity of the PCV derived secondary sprout (Figure 3N).

We identified that this differential movement is largely specified in the ECs of the individual ISV but can be reverted by flow-induced directional movement to obtain a balanced number of arteries and veins (Figure 4). Blocking the latter by inhibiting blood flow showed that specification alone leads to a 60/40 ratio of arteries and veins (Figure 2A), indicating that flow-mediated compensation induces re-specification of a subset of arteries into veins even in wild-type embryos. A similar ratio was observed in artery-*versus* vein-specific primary ISV EC behavior when blocking secondary sprouting (Figure 3J). The extent of the compensation effect is adaptable to compensate for larger deviations from the optimal 50/50 ratio as shown by the mosaic Notch activation experiments. The molecular basis of the newly identified early aISV and vISV specification, remains to be determined. Our data identify that Notch signaling, a long known arterial determinant appears to specify arterial identity in the zebrafish trunk vasculature, through instructing endothelial polarity and movement. Further studies will need to uncover how flow can overwrite the early ISV specification. Previous work suggests that it could do so by directly modulating Notch levels [21], although the possibility of a distinct specification pathway cannot be excluded

**Figure 4.**
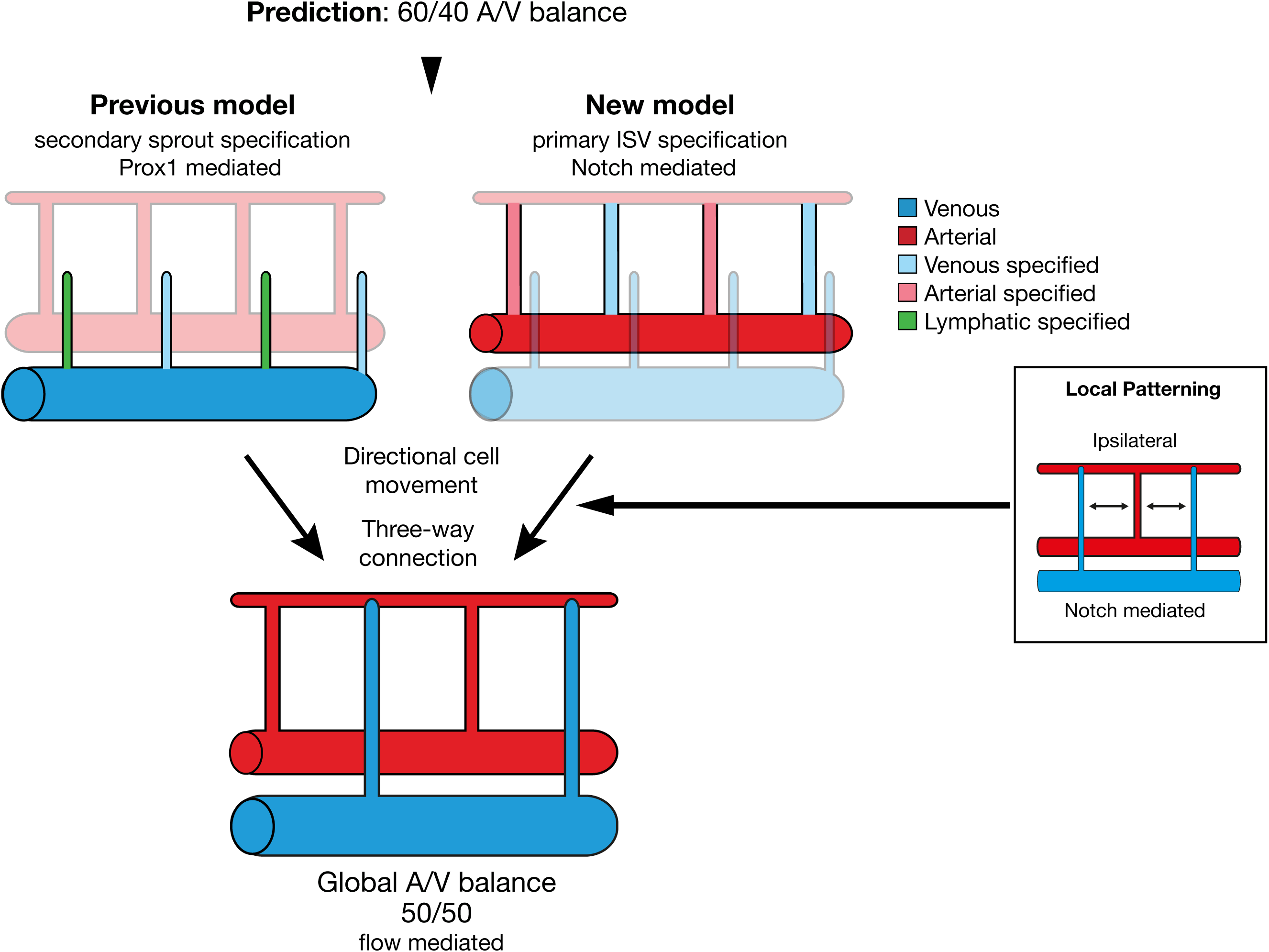
Alternative mechanism of arterio-venous specification in the zebrafish trunk.

In conclusion, our work identifies transient arterio-venous connections as intermediate structures that resolve through a combination of pre-specified and flow-induced directional migration of ECs. We propose that vascular remodeling, based on a dual mechanism of molecular specification and flow-mediated patterning control, provides the necessary plasticity to allow formation of an overall balanced and efficiently perfused vascular network in the zebrafish trunk.

## METHODS

### Zebrafish husbandry and transgenic lines

Zebrafish (*Danio rerio*) were raised and staged as previously described [22]. The following transgenic lines were used: *Tg[fli1a:GFP]*^*y1*^[9], *Tg[fli1a:nEGFP]*^*y7*^ [23], *Tg[fli1a:dsRedEX]*^*um13*^[24], *Tg[gata1a:dsRed]*^*sd2*^[25], *TgBAC[prox1:KalT4-UAS:uncTagRFP]*^*nim5*^ [26, 27], *Tg[-0.8flt1:RFP]*^*hu5333*^ [3], *Tg[fli1a:B4GalT1-mCherry]*^*bns9*^ [28], T*g[fli1a:pecam1-EGFP]*^*ncv27*^ [29]. For growing and breeding of transgenic lines we comply with regulations of the ethical commission animal science of KU Leuven and MDC Berlin.

### Vessel patterning analysis

Figures 1B-C, 1I, 2A-B, S1A-B show analyses of the regularity of local and global vessel arrangements of aISVs and vISVs in the zebrafish trunk. To perform those analyses, 6dpf zebrafish embryos were screened under a fluorescent stereomicroscope (Leica M205 FA) to identify the sequence of arterial and venous ISVs on both flanks of a 10-somite segment in the trunk, starting after the junction of DA and PCV (i.e. somites 5-15). The identity of arteries and veins was determined by their connection to respectively the DA or the PCV, and by direction of blood flow. This manual characterization yielded for each vessel its ISV type, its collateral neighbor’s ISV type, as well as the left ISV neighbor type for all but the leftmost vessels, and the right ISV neighbor type for all but the rightmost vessels. In mosaic NICD overexpression experiments, in addition to the vessel type it was also registered if a given vessel contained a NICD overexpressing cell (used in Figure 1I). This dataset was then analyzed to obtain the global and local distribution patterns of aISVs and vISVs for different experimental conditions using a custom-made Python script.

To obtain the global vessel ratio, the number of aISVs and vISVs of all ISVs was compared in Figures 1A,G,H and Figure 2A,C.

To study the local ISV patterning, the relative frequency of aISVs given the type of the neighboring vessels was obtained: Figure 1C shows the frequency of aISVs given the type of the contralateral neighbor (aISV or vISV); Figures 1B, 1I and 2B show the frequency of aISVs given the vessels’ corresponding number of ipsilateral aISV neighbors (0, 1 or 2 aISVs); Supplementary Figure 1A shows the frequency of aISVs for the given number of aISVs in the 2-neighborhood of the vessel (0, 1, 2, 3 or 4 aISVs); and Supplementary Figure 1B shows the frequency of aISVs given the type of the right neighbor (aISV or vISV).

### Mosaic overexpression using Tol2 transgenesis

Transgenic zebrafish embryos *Tg[fli1a:eGFP]*^*y1*^ were injected at one-cell stage with 100 pg of Tol2 mRNA [30] and 15 pg of plasmid DNA pTol2Fliep-N1aICD-basfli-mCherry [31]. Embryos were raised at 28°C and screened for transient expression at ˜30 hpf.

Quantification of arterial and venous ISV distribution was performed in 6 dpf embryos by scoring their percentile presence in 10 consecutive somite segments in the trunk after the junction of DA and PCV (i.e. somites 5-15).

### Live imaging

Embryos were anaesthetized in 0,014% tricaine (MS-222, Sigma), mounted in a 35 mm glass bottom petri dish (0.17 mm, MatTek) using 0,6% low melting point agarose (Sigma) containing 0,014% tricaine, and bathed in Danieau’s buffer containing 0,007% tricaine and 0,003% PTU. Time-lapse imaging was performed using a Leica TCS SP8 upright microscope with a Leica HCX IRAPO L x25/0.95 water-dipping objective and heating chamber, or on an upright 3i spinning-disc confocal using Zeiss Plan-Apochromat ×63/1.0 NA, x40/1.0 NA or x20/1.0 NA water-dipping objectives. Image processing was performed using Fiji software [32].

### Tricaine treatment

To slow down heart rate during the secondary sprouting and ISV remodeling process embryos were treated with 0,03% (2x) tricaine (MS-222, Sigma) between 31 and 52 hpf, after which the compound was washed away again.

### Cell polarity analysis

To analyze polarity of ECs during vascular remodeling, time-lapse movies were made of transgenic *Tg[fli1a:GFP]*^*y1*^*; Tg[fli1a:B4GalT1-mCherry]*^*bns9*^ embryos during vascular remodeling in the trunk (˜32 hpf to ˜54 hpf). Polarity arrows from the center of the nucleus to the center of the golgi complex were drawn manually using Fiji software. For every primary ISV cell the polarity was scored per time point: dorsal polarity, ventral polarity or unpolarized, depending on the relative position of golgi and nucleus, i.e. respectively, golgi dorsal, ventral or parallel to the nucleus in respect to the local angle of the ISV. Per developmental phase all scores were added over the different timepoints (phase I: 1 to 2.5 hours before secondary sprout connection; phase II: during three-way connection; phase III: 1 to 2.5 hours after resolution of the three-way connection).

### Cell movement analysis

To analyze the upward movement of ECs within ISVs of *Tg[fli1a:GFP]*^*y1*^ or *Tg[fli1a:nEGFP]*^*y7*^*; Tg[fli1a:dsRedEX]*^*um13*^ embryos in different stages of development, we use a mix of manual segmentation of developmental timelapse and computational analysis in Python (Figures 3F & 3M). Confocal stacks were registered using StackReg (ImageJ plugin - http://bigwww.epfl.ch/thevenaz/stackreg/). The cells’ distance to the DA was tracked through time in a 2D maximum projection manually in Fiji, and combined with the information about the later fate (aISV or vISV) of the vessel containing the cell, and the current phase of development of the ISV (cf. Figure 3A). To obtain the *upward speed* of the cells in the individual phases of the vessel development, the initial distance of the cell’s nucleus to the aorta was compared to the final position in the given phase (in [μm]) divided by the duration of the cell’s trajectory in the phase (in [min]). Note that in this definition a positive *upward speed* corresponds to an increasing distance away from the DA (i.e. movement towards the dorsal side of the trunk), while a negative *upward speed* corresponds to an average movement towards the DA.

### Morpholino knockdown

Morpholinos against *ccbe1* and *dll4* were used as previously described and injected at 5ng and 10ng/embryos respectively [17, 18].

### In vitro flow chamber experiments

HUVECs (Promocell, primary cells from pooled donors) were seeded and grown to confluency on a slide in EBM2 medium (Promocell) coated with gelatin. Unidirectional laminar shear stress was applied to confluent HUVECs using a parallel plate chamber system [33] for 24 hours and cells were treated with 5μM DAPT or a similar amount of DMSO in controls for the duration of the experiment. Local shear stress was calculated using Poiseuille law and averaged 20 dyn/cm^2^. Cells were fixed in 100% methanol for 10min at -20°C and stained for DAPI, VE-cadherin (Santa Cruz, sc-6458, dilution 1/100) and GM130 (BD Biosciences, 610822, dilution 1/400) [34]. Matlab was used to analyze cell orientation (direction of the main axis of the nucleus) and cell polarity (angle between center of the nucleus and center of the golgi).

### Statistical Analyses

No statistical method was used to predetermine sample size.

Data represent mean ± SEM of representative experiments. Statistical tests were conducted using Prism (GraphPad) software. Adequate tests were chosen according to the data to fulfil test assumptions. Sample sizes, number of repeat experiments, performed tests and p-values are indicated per experiment in Supplemental Table 1. The angle repartitions of the flow chamber experiments were analyzed using Kuiper two-sample test, a circular analogue of the Kolmogorov-Smirnov test. A p-value < 0.05 was considered statistically significant.

Zebrafish embryos were selected on the following pre-established criteria: normal morphology, beating heart, presence of circulating red blood cells. The experiments were not randomized. For every experiment treated and control embryos were derived from the same egg lay. The investigators were not blinded to allocation during experiments and outcome assessment.

### Data availability statement

The data that support the findings of this study are available from the corresponding author upon reasonable requests.

### Code availability statement

The codes that were used for computational analysis of *in vivo* cell movement (Python) and *in vitro* cell orientation (Matlab) are available from the corresponding author upon reasonable request

---

**Supplementary Figure 1**

A) Ipsilateral neighborhood analysis of vessel identity with 4 neighbors in 6dpf WT embryos (n=3 experiments, 74 embryos, 888 ISVs).

B) Ipsilateral neighborhood analysis of vessel identity with 1 neighbor in 6dpf WT embryos (n=3 experiments, 74 embryos, 1332 ISVs).

C) Stills from time-lapse movie (Supplementary Movie S2) in *Tg[fli1a:GFP]*^*y1*^*;Tg[gata1a:DsRed]*^*sd2*^ labeling ECs in green and blood cells in red showing formation of a transient perfused three-way connection (C’-C”) as circulating blood cells can be observed in the DA-ISV-secondary sprout-PCV shunt. In panel C”’ it is clear that the ISV-PCV connection is disconnected again and the secondary sprout takes part in lymphatic development, whereas strong dorsal flow is established in the arterial ISV.

D) Stills from time-lapse movie (Supplementary Movie S3) in *Tg[fli1a:GFP]*^*y1*^ embryos mosaically overexpressing a pT2-zN1aICD-basfli-mCherry construct showing formation of a transient perfused three-way connection between a wild-type secondary sprout and a NICD overexpressing (NICD^OE^) primary ISV, indicating that Notch activation does not prevent interaction between primary ISVs and secondary sprouts.

Scale bars, 10μm

**Supplementary Figure 2**

A) Example of two different type of regression behavior, imaged in T*g[fli1a:pecam1-EGFP]*^*ncv27*^*; Tg[-0.8flt1:RFP]*^*hu5333*^ 5ng *MO-ccbe1* embryos: Full disconnection and thin membrane connection left.

B) Quantification of the nature of ISV regression behavior in *MO-ccbe1* embryos (n=4 experiments, 37 embryos).

C) Quantification of the nature of ISV regression behavior in *MO-ccbe1*/*MO-dll4* embryos (n=4 experiments, 47 embryos).

D) Quantification of EC polarity in aISVs and vISVs of *Tg[fli1a:GFP]*^*y1*^*;Tg[fli1a:B4GalT-mCherry]*^*bns9*^ *MO-dll4* and WT embryos at 3 different time points: I) 2.5 hours before secondary sprout connection, II) during three-way connection and III) 2.5 hours after resolution (n=7 WT aISV, 8 WT vISV, 10 *MO-dll4* vISV).

E) Flow chamber experiment. A confluent layer of HUVEC cells was exposed to high shear stress and treated with 5μM DAPT or DMSO (control). Cells were stained for DAPI (nuclei), VE-cadherin (cell boundaries) and GM130 (golgi). Cell orientation was analyzed by plotting the direction of the main axis of the nucleus. Cell polarity was determined by defining the angle between the center of the nucleus and the center of the Golgi. The graphs represent the percentage of cells with a certain angle of direction or polarity relative to the direction of flow (10° intervals), the red line indicates the mean.

## Supplementary Movie Legends

(1) Time-lapse imaging of a *Tg[fli1a:EGFP]*^*y1*^ */ Tg[-0.8flt1:RFP]*^*hu5333*^ embryo showing ISV remodeling into an aISV and a vISV from 26 to 55 hpf (frame interval: 15 minutes). In both cases, a lumenized connection is formed between the secondary sprout and the primary ISV. In the case of the formation of an aISV, the connection is lost again and the secondary sprouts forms lymphatic precursors at the horizontal myoseptum (Parachordal lymphangioblasts). In case of vISV remodeling, the secondary sprout connection is stabilized and the connection between primary ISV and DA regresses. Scale bars, 50μm

(2) Time-lapse recording of a *Tg[fli1a:GFP]*^*y1*^*;Tg[gata1a:DsRed]*^*sd2*^ embryo between 35 and 40 hpf (frame interval: 2 minutes) showing a transient perfused three-way connection in which the ISV-PCV connection dissociates again to form an arterial ISV and a lymphangiogenic sprout. Scale bars, 10μm

(3) Time-lapse recording of a *Tg[fli1a:GFP]*^*y1*^ embryo mosaically overexpressing a pT2-zN1aICD-basfli-mCherry construct between 31 and 43 hpf (frame interval: 6 minutes) showing formation of a transient perfused three-way connection between a wild-type secondary sprout and a NICD^OE^ primary ISV. Scale bars, 15μm

(4) Time-lapse imaging of *a Tg[fli1a:GFP]*^*y1*^*;Tg[fli1a:B4GalT-mCherry]*^*bns9*^ embryo showing EC polarity in an arterial ISV from 32 to 51 hpf (frame interval: 15 minutes). Scale bars, 25μm

(5) Time-lapse imaging of *a Tg[fli1a:GFP]*^*y1*^*;Tg[fli1a:B4GalT-mCherry]*^*bns9*^ embryo showing EC polarity in a venous ISV from 33 to 55 hpf (frame interval: 15 minutes). Scale bars, 25μm

(6) Time-lapse imaging of *Tg[fli1a:pecam1-EGFP]*^*ncv27*^, *Tg[-0.8flt1:RFP]*^*hu5333*^ embryo from 29 to 53 hpf showing ISV remodeling into aISVs and vISVs (frame interval: 15 minutes). Scale bars, 50μm

(7) Time-lapse imaging of a T*g[fli1a:pecam1-EGFP]*^*ncv27*^ *Tg[-0.8flt1:RFP]*^*hu5333*^ *MO-ccbe1* embryo from 30 to 54:30 hpf showing ISV regression in the absence of secondary sprouting (frame interval: 15 minutes). Scale bars, 50μm

(8) Time-lapse imaging of a *Tg[fli1a:EGFP]*^*y1*^*/ Tg[-0.8flt1:RFP]*^*hu5333*^ *MO-ccbe1/MO-dll4* embryo from 30 to 54h30 hpf showing ISV regression in the absence of secondary sprouting (frame interval: 15 minutes). Scale bars, 50μm

## AUTHOR CONTRIBUTIONS

I.G., B.C., S.A., V.G., A.-C.V. and H.G. designed the experiments. I.G., B.C., S.A., V.G., A.R. and A.-C.V. performed the experiments and analyzed the data. I.G., B.C., S.A. and H.G. wrote the manuscript.

